# PRDM3 and PRDM16 define cranial neural crest cell states in zebrafish development

**DOI:** 10.64898/2026.05.14.725231

**Authors:** Lomeli C. Shull, Silvia Meyer-Nava, Bryanna Saxton, Qootsvenma Denipah-Cook, Fahmida Raha, Julaine Roffers-Agarwal, Job Flores, Ezra Lencer, Srinivas Ramachandran, Kristin B. Artinger

## Abstract

Cartilage and bone that comprise craniofacial structures as well as neurons and glia of the peripheral nervous system are derived from a multipotent population of cranial neural crest cells, that respond to both cell intrinsic and extrinsic cues to differentiate into precise cell states. Both a genetic and epigenetic regulatory network are required for each step in the differentiation process, involving transcription factors, histone modifiers and chromatin remodelers. Here, we examined the direct transcriptional targets of two histone methyltransferases, Prdm3 and Prdm16 in zebrafish neural crest cells at 48 hours post fertilization in zebrafish. Using CUT&RUN, we examined both direct DNA binding and nucleosome association. At this stage of development, CUT&RUN fragment size analysis indicated that Prdm3 and Prdm16 are largely associated with nucleosomes. We further analyzed these nucleosome peak sets to identify 6 clusters where differential binding of Prdm3 and Prdm16 and differential enrichment of gene ontology terms for target genes was observed. We validated gene expression in each cluster by *in situ* hybridization chain reaction (HCR) at 48 hpf demonstrating that *prdm3* and *prdm16* mutants exhibit corresponding changes in gene expression of the putative gene targets identified. Finally, we performed CUT&RUN-qPCR in *prdm3* and *prdm16* mutant zebrafish embryos and demonstrated reduced binding at putative target loci. Together these data suggest that Prdm3 and Prdm16 regulate their transcriptional targets primarily by binding nucleosomes around their putative target loci to control downstream gene expression.

**Highlights:** Prdm3 and Prdm16 associate with nucleosomes for regulation of gene expression

Gene targets are altered in prdm3 and prdm16 mutant zebrafish

Reduced binding is observed in respective mutants

## Introduction

During craniofacial development, multipotent cranial neural crest cells (cNCC) differentiate into precise cell states that make them unique from other cells in the head (Bronner and LeDouarin (2012); (Le Douarin, 1982). This is because cNCCs give rise to a diverse array of cell types, including chondrocytes and osteoblasts that will form the cartilage and bone of the facial structures, pigment cells, and neurons and glia of the peripheral nervous system among other derivatives (Minoux and Rijli, 2010; Selleri and Rijli, 2023). The gene regulatory networks (GRN) and signaling pathways orchestrating these differentiation processes must be tightly controlled, as alterations at any step contributes to the etiology of congenital birth defects affecting the formation of the craniofacial complex (Simoes-Costa and Bronner, 2015) (Martik and Bronner, 2021; Mitani et al., 2006).

Neural crest cell (NCC) lineage trajectories have been the focus of many more recent studies, due to the precise temporal and spatial order of specification and differentiation. It is now understood that a highly regulated NCC GRN is required to define different cell fates. The unique cell states upregulate the expression of specific transcription factors in response to signals including BMPs, Wnts, and FGFs as well as others. cNCCs are first specified distinctly from the neural plate and the non-neural ectoderm, defining the neural plate boarder. Here, NCCs express neural crest specifier genes including *sox10*, *foxd3* and *snail2*. Following specification, cranial NCCs undergo an epithelial-to-mesenchymal transition (EMT) and migrate into the pharyngeal arches, where they begin to differentiate into chondrocytes and bone by upregulating *col2a1* and *runx2,* respectively in response to endothelin signaling. cNCCs also intermingle with ectodermal placode cells and migrate to areas of the cranial ganglia to form neurons and glia (Schlosser, 2005). They respond to signals including Wnt and Shh and upregulate the expression of neuronal markers *elav3, isl1* and *neurog1* and glial markers such as *foxd3* (Kurosaka et al., 2015).

Epigenetic regulators direct nucleosome dynamics and accessibility of transcriptional machinery to the DNA by post-translationally modifying histones and also altering nucleosome stability, which results in precise temporal and spatial control of gene expression. There is increasing experimental evidence that epigenetic factors play an equally important role in NCC fate decisions and all stages of cranial neural crest development (Shull and Artinger, 2024) (Selleri and Rijli, 2023). Many of these modifiers are also mutated in human disease. For example, loss of the ATP-dependent chromatin remodeler CHD7 in *Xenopus*, zebrafish and mice results in craniofacial defects, similar to the phenotypes of human patients with CHARGE syndrome that carry mutations in CHD7 (Bajpai et al., 2010; Hurd et al., 2010; Lalani et al., 2006; Layman et al., 2010; Sperry et al., 2014). Similarly, human mutations in KAT6A, an acetyltransferase of H3K9 that activates gene expression, results in Arboleda-Tham Syndrome in which patients have dysmorphic facial features. These disease phenotypes have been replicated using animal models including mice and zebrafish (Vanyai et al., 2019; Voss et al., 2012).

Two histone methyltransferases belonging to the PRDM family, PRDM3 (MECOM/EVI1) and PRDM16, have been identified through human GWAS studies to affect normal human facial morphology (Shaffer et al., 2016). Additionally, SNPs in these genes are associated with cleft lip with or without cleft palate (Jugessur et al., 2010; Li et al., 2019; Shaffer et al., 2016). These two paralogs are heavily involved in lineage decisions during development of many tissues and organ systems. Both factors methylate lysine residues on H3K9 and H3K4 through their intrinsic methyltransferase activity and/or bind DNA directly via zinc finger domains to regulate downstream gene expression, although endogenous enzymatic activity has not been definitively shown *in vivo*. Prdm3 is important for mediating differentiation programs for hematopoietic cell populations, while Prdm16 controls the cell fate switch between muscle and brown fat (Kajimura et al., 2008; Kataoka et al., 2011; Sato et al., 2008; Seale et al., 2008). Both have recently been shown to control lung cell fate determination (He et al., 2024). In neural crest development, Prdm3 and Prdm16 are strongly expressed within the craniofacial complex and importantly control zebrafish and mammalian craniofacial chondrocyte maturation and differentiation. In mice, *Prdm3* null mutant embryos are embryonic lethal while conditional ablation in the neural crest under the control of the Wnt1-Cre driver results in Meckel’s cartilage chondrocyte disorganization suggesting defects in chondrocyte differentiation (Shull et al., 2022; Shull et al., 2020). *Prdm16* null animals and neural crest-specific conditional ablation develop palate defects, abnormal chondrocyte differentiation, and mandibular hypoplasia (Bjork et al., 2010; Shull et al., 2022; Shull et al., 2020; Warner et al., 2013). Further, we have previously shown in zebrafish that *prdm3* and *prdm16* paralogs function to promote cartilage differentiation by spatiotemporally balancing Wnt/β-catenin activity, both at the transcriptional level and epigenetically by controlling chromatin accessibility of downstream Wnt/β-catenin targets (Shull and Artinger, 2024; Shull et al., 2022). However, their direct transcriptional and chromatin targets in neural crest development remain largely unknown.

Here, we performed CUT&RUN in neural crest cells to further define the direct transcriptional targets of Prdm3 and Prdm16. By analyzing CUT&RUN fragment size distribution we assessed modes of transcription factor activity: direct DNA binding indicated by small DNA fragments vs nucleosome association indicated by larger fragments (Skene and Henikoff, 2017; Tonsager et al., 2025; Trouth et al., 2024). Nucleosome-sized fragments yielded the vast majority of peaks in the CUT&RUN datasets, pointing to nucleosome association by Prdm3 and Prdm16. Furthermore, based on enrichment of Prdm3, Prdm16, and H3K27ac, we were able to identify 6 clusters of peaks. The genes associated with the peaks showed differential enrichment of gene ontology (GO) terms and were predicted to be regulated by either Prdm3 or Prdm16 or by both. To determine if patterns of Prdm3/Prdm16 binding defined how target genes were regulated, we examined gene expression for representative genes in each cluster by HCR in situ hybridization at 48 hpf demonstrating that *prdm3* and *prdm16* mutants have changes in gene expression of the targets identified based on Prdm3 and Prdm16. Finally, we performed CUT&RUN qPCR in *prdm3* and *prdm16* mutant zebrafish embryos and identified binding of Prdm3 and Prdm16 that is consistent for the targets based on the cluster the targets belonged to.

## Methods

### Zebrafish husbandry

Zebrafish were maintained as previously described (Westerfield, 2000). The wildtype (WT) strain used was AB (ZIRC) and the mutant lines used were *prdm3^co1013^(*referred to as *prdm3^-/-^)* and *prdm16^co1027^ (*referred to as *prdm16^-/-^)*. All experiments were reviewed and approved by the Institutional Animal Care and Use Committee (IACUC) at the University of Minnesota (protocol # 2210-40517A) and University of New Mexico (protocol #24-201487-B-MC and #24-201481-MC) and conform to the NIH regulatory standards of care and treatment. Zebrafish lines can be obtained from the lead contact.

### FAC-Sorting of Neural Crest Cells

At 48 hpf, Tg(*sox10*:mRFP)-positive embryos were stage-matched and selected under a fluorescence dissecting microscope. To prepare single-cell suspensions for FACS, ∼1000 embryos were dechorionated and rinsed in 1XDPBS (Calcium and Magnesium free). Embryos were dissociated in Accumax (Innovative Cell Technologies, AM-105) with DNaseI (Roche Diagnostics). Samples were incubated at 31°C and gently agitated with pipetting every 10 minutes for 1 hour. Following the digest, a wash solution containing 1xDPBS and DNaseI was added to stop the dissociation. Cells were filtered through a 70 µM nylon mesh strainer, centrifuged at 2000 rpm for 5 minutes at 4°C and resuspended in FACS basic sorting buffer (1 mM EDTA, 25 mM HEPES (pH 7.0), 1% fetal bovine serum in 1xDPBS). The single cell suspension was stained with DAPI at a dilution of 1:10000 and kept on ice. RFP positive cells were collected on a MoFlo XDP100 cell sorter (Beckman-Coulter) with a 100 um nozzle tip and collected in 1xDPBS. After FACS, RFP-positive cells were subjected to CUT&RUN.

### CUT&RUN

CUT&RUN was performed on FAC-Sorted Tg(*sox10*:mRFP)-positive neural crest cells isolated from 48 hpf zebrafish. Briefly, following FACS, approximately 500,000 cells were incubated on activated Concanavalin A conjugated paramagnetic beads (Epicypher) for 10 minutes at room temperature. Cell-bound beads were resuspended in antibody buffer (20 mM HEPES, pH 7.5; 150 mM NaCl; 0.5 mM Spermidine (Invitrogen); 1x Complete-Mini Protease Inhibitor tablet (Roche Diagnostics); 0.01% Digitonin (Sigma Aldrich); 2 mM EDTA) and incubated in antibodies directed toward Prdm3 (Aviva), Prdm16 (Novus Biologics), H3K27ac (Cell Signaling), or IgG (Epicypher) overnight with rotation at 4°C. The following day, cells were washed in Digitonin Buffer wash solution (20 mM HEPES, pH 7.5; 150 mM NaCl; 0.5 mM Spermidine (Sigma Aldrich); 1x Complete-Mini Protease Inhibitor tablet (Roche Diagnostics); 0.01% Digitonin (Sigma Aldrich) twice and then incubated with pAG-MNase (EpiCypher) for 10 minutes at room temperature. After another two washes in Digitonin Buffer, 1 ul of 100 mM Calcium Chloride was added to each sample. Samples were then incubated at 4°C for 2 hours with rotation. The digestion reaction was stopped with the addition of Stop Buffer (340 mM NaCl, 20 mM EDTA, 4 mM EGTA, 50 ug/ml RNase A and 50 ug/ml glycogen) for 10 minutes at 37°C. DNA fragments were eluted and purified using a DNA Clean and Concentrator Kit (Zymo Research). For sequencing, libraries were prepared from purified CUT&RUN fragmented DNA using the NEBNext Ultra II DNA Library Prep Kit for Illumina (New England Biolabs) as per the manufacturer’s instructions. Amplification of the libraries was performed according to the guidelines outlined by EpiCypher: 98°C for 45s; 98°C for 15s and 60°C for 10s, 14 cycles; and 72°C for 1 minute. Samples were then subjected to paired-end 150 bp sequencing on the Illumina NovaSEQ 6000 system at Novogene Corporation (Sacramento, CA, USA). CUT&RUN experiments were performed in duplicate for two biological replicates.

### Bioinformatics Analysis

Libraries were sequenced on an Illumina NovaSeq platform PE 2x150bp, to a depth between 30-50 million PE reads. Adapters were trimmed from raw fastq files using Cutadapt v3.5 (Martin, 2011) (-a AGATCGGAAGAGCACACGTCTGAACTCCAGTCAC -A GATCGTCGGACTGTAGAACTCTGAACGTGTAGATCTCGGTGGTCGCCGTATCATT -- minimum-length=25). Trimmed reads were aligned to the zebrafish genome (GRCz11) with bowtie2 v2.4.5 (--end-to-end --very-sensitive)(Langmead and Salzberg, 2012). Samtools (Li et al., 2009) and bedtools (Quinlan and Hall, 2010) were used for processing aligned reads from sam to bed files. Coverage at 10 bp windows genome-wide was calculated as the number of short and long reads that mapped at that window, normalized by the factor N: N = 1345101833/(Total Number of Mapped Reads)

Short reads were defined as reads between 30 and 80 bp, whereas long reads (“nucleosomal”) were defined as reads between 120 and 200 bp. We called peaks as previously described(Rao et al., 2022), using a custom Python script (https://github.com/satyanarayan-rao/tf_nucleosome_dynamics/blob/main/CUTnRUN_Peak_Calling/call_peaks_iterative.py).

Briefly, the peak-caller smoothes the 10-bp coverage tracks defined above with a Savitzky-Golay filter (Savitzky and Golay, 1964) available as a SciPy (Harris et al., 2020) method “signal.savgol_filter” with parameters window_length = 9, polyorder = 1. We determined the cutoff for each dataset by iteratively eliminating outliers and used the “find_peaks” method in SciPy to call peaks that were separated by at least 250 bp. The peaks were then filtered to retain only those with at least 4-fold higher enrichment with a given epitope compared to IgG.

We focused on peaks called using long fragments for further analysis (explained in Results). The peaks across Prdm3, Prdm16, and H3K27ac were combined into a single list and the log2 fold change of CUT&RUN enrichment over IgG across the three datasets was calculated. The log2 fold change for Prdm3, Prdm16, and H3K27ac were used to perform k-means clustering of the peaks with k=6. Genes were assigned to peaks using the program GREAT(McLean et al., 2010). Published ATAC-seq datasets were processed similarly to CUT&RUN data for generating scores at peaks. Metaplots and heatmaps were generated for CUT&RUN and ATAC-seq datasets using deepTools (Ramirez et al., 2016).

### CUT&RUN-qPCR

Prior to CUT&RUN paired with qPCR on mutant zebrafish, 48 hpf embryos from each strain were genotyped. To do this, embryos were placed in a low dose of tricaine (MS-22, Sigma Aldrich) and tail clips were collected from individual embryos and placed directly into Tissue Phire Direct PCR Master Mix or samples were incubated in lysis buffer [10 mM Tris-HCl (pH 8.0), 50 mM KCl, 0.3% Tween-20, 0.3% NP-40, 1 mM EDTA] for 10 min at 95°C. Lysates were then digested with 50 µg Proteinase K at 55°C for 2 h, followed by Proteinase K inactivation at 95°C for 10 min with primers specific to each mutant strain (for *prdm3*: (F) 5’-CTGTGGGCAGATGTTTAGCA-3’, (R) 5’-ACTATGATGCCGGTTTGTCC-3’; for *prdm16*: (F) 5’-TAAGCAATTATGTGATGCCGTC-3’, (R) 5’-CTTTTCACAGTCTTTGCACTCG-3’). Following PCR cycling, products were run out on 3-4% agarose gels. Once confirmed, heads of wildtype and each respective mutant were cut off and pooled (about 20-30 animals per genotype).

Tissue dissociation in preparation for CUT&RUN was performed as described above. The same number of head were used and cells were counted to ensure the same number of cells were used in each CUT&RUN reaction for each genotype. CUT&RUN was performed on the cells as described above. CUT&RUN eluted fragmented DNA was amplified using NEBNext UltraII DNA Library prep kit as per manufacturer’s instructions. RT-PCR was performed with primers designed to flank putative Prdm3, Prdm16 or H3K27ac binding motifs at the promoter region of targets or negative IgG control. The fold enrichment for the abundance of Prdm3, Prdm16 or H3K27ac of the amplified regions was normalized to IgG controls across three experiments and three technical replicates.

### Hybridization Chain Reaction (HCR) V3.0

All HCR probes (*snail1a, foxi1,myf5, foxd3, neurog1, mkxa, prickle1a, sox2, pax3a, isl1a, chd5*) were purchased from Molecular Instruments (www.molecularinstruments.com). Whole mount HCR was performed according to the manufacturer’s instructions (Choi et al., 2016; Choi et al., 2018) and as previously described (Truong et al., 2023). Briefly, embryos were fixed overnight at 4°C in 4% PFA, washed in PBS, and dehydrated and permeabilized in two 10-minute washes in 100% MeOH at room temperature. Embryos were stored for at least 24 hours at -20°C in fresh MeOH. A graded series of MeOH/PBST solutions was used to rehydrate the embryos (75%, 50%, 25%, 0%). Embryos were then treated with proteinase K (10 μg/mL) for 15 minutes (48 hpf), washed twice in PBST, fixed for 20 minutes in 4% PFA, and then washed five times in PBST. Hairpins were heated to 95°C for 90 seconds and cooled. Embryos were stored in PBS at 4°C protected from light. Whole embryos were mounted in 0.2% low melt agarose and imaged on Zeiss LSM confocal. Embryos were then genotyped following Shull et al (2022) with slight modifications. Following DNA extraction, PCR was performed in M buffer (2mM MgCl_2_, 13.7 mM Tris-HCl [pH 8.4], 68.4mM KCl, 0.001% gelatin, 1.8 mg/mL protease-free BSA, 136 μM each dATP/CTP/GTP/TTP) with GoTaq Flexi (Promega) and genotyping primers (for *prdm3*: (F) 5’-CTGTGGGCAGATGTTTAGCA-3’, (R) 5’-ACTATGATGCCGGTTTGTCC-3’; for *prdm16*: (F) 5’- TAAGCAATTATGTGATGCCGTC-3’, (R) 5’-CTTTTCACAGTCTTTGCACTCG-3’) and products were resolved on a 4% agarose gel. Quantification of signal intensity was performed by creating a ROI around the pharyngeal arches using sox10+ cells, measuring the mean intensity as compared to background and calculating the corrected total cell fluorescence (CTCF) per tissue area of the pharyngeal arches.

## Results

### Prdm3 and Prdm16 bind both similar and unique sites in the genome

To determine the chromatin binding of Prdm3 and Prdm16, we performed CUT&RUN on FAC sorted *Tg(sox10:mRFP)* NCCs from zebrafish embryos at 48 hpf, using antibodies against Prdm3, Prdm16, H3K27ac and IgG as a control. From the CUT&RUN sequencing data, we determined genome-wide distribution of short fragments (<80 bp) and nucleosomal fragments (120-200 bp) and identified peaks. The number of peaks obtained from the distribution of short fragments were 606, 572, and 941 respectively for Prdm3, Prdm16, and H3K27ac. Nucleosomal peaks were much higher in number: 36,647, 26,863, and 57,688 respectively for Prdm3, Prdm16, and H3K27ac (**Table 1**). These results suggest that while there might be a small fraction of sites on the genome where Prdm3 and Prdm16 bind directly to DNA, they preferentially associate with nucleosomes to remodel the chromatin landscape at this developmental timepoint (Rao et al., 2022) (**Figure 1A-C**). Hence, we focus on the nucleosome-associated peak for the rest of the manuscript. We combined peaks from Prdm3, Prdm16, and H3K27ac datasets and calculated the CUT&RUN enrichment across the three datasets from the combined list of peaks. We then performed k-means cluster (with k=6) to identify patterns of binding across Prdm3, Prdm16 and H3K27ac relative to IgG. Heat map of the log2 fold change over IgG ordered by the 6 clusters show the different patterns of overlap between the three targets (**Figure1D, E**). Cluster 1 consists of peaks with high enrichment primarily in H3K27ac, cluster 2 consists of peaks with high enrichment in H3K27ac and Prdm3, and cluster 3 consists of peaks with high enrichment in Prdm16 and H3K27ac. Cluster 4 shows highest enrichment for Prdm16, with significant enrichment for Prdm3, and H3K27ac as well. Cluster 5 shows high enrichment only for Prdm3. Cluster 6 consists of a small fraction of sites with enrichment for only Prdm3 and Prdm16, but not H3K27ac (**Figure1D, E**). *k*-means clustering revealed 6 different clusters with unique enrichment patterns, indicative of unique genomic binding patterns (**Figure 1E**), as demonstrated with metaplots for each cluster (**Figure 2A-E**). Some bound targets were unique to Prdm3 and Prdm16 respectively, and other targets have both bound. These patterns also reflect differences in the presence of H3K27ac within each cluster, for example, Cluster 1 (CL1) has high H3K27ac at both the promoter and along histones.

**Figure 1:**
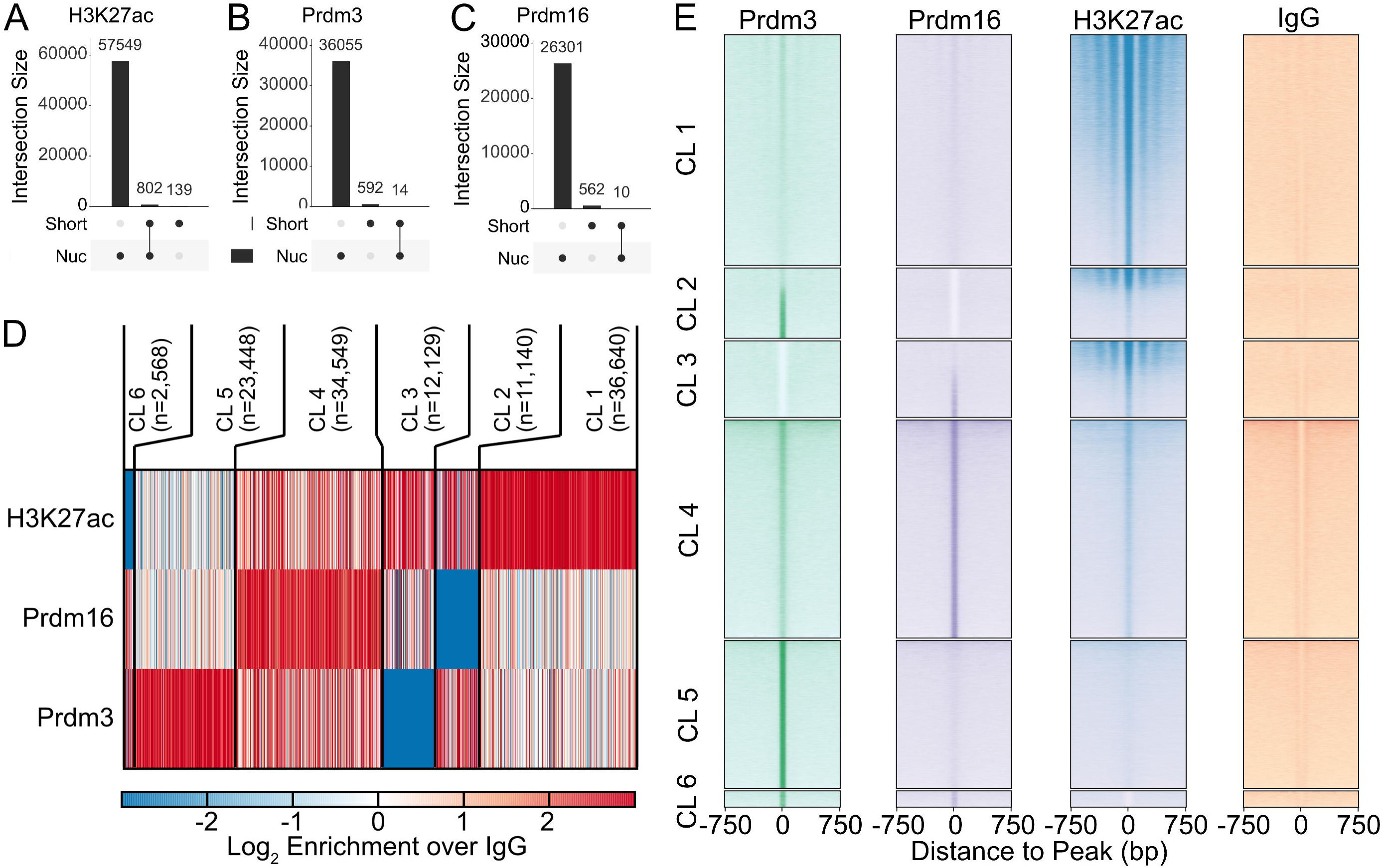
CUT&RUN identifies unique targets of Prdm3 and Prdm16. CUT&RUN performed on sox10+ sorted cells at 48 hpf for Prdm3, Prdm16 H3K27ac, and IgG. (A-C) UpSet plots for overlap of peaks called from short fragment enrichment and long fragment enrichment in H3K27ac, Prdm3, and Prdm16 demonstrates that the majority of the enriched peaks for Prdm3 (36055) and Prdm16 (26301) are nucleosome associated peaks, similar to H3K27 (57549). Few are short read peaks which suggest direct DNA binding (D) Heat map of log2 enrichment over IgG plotted for Prdm3, Prdm16 and H3K27ac. The peaks are ordered based on the cluster (CL) they belong to and show the number of peaks total in each cluster. The majority are in CL1 and CL4. CL represents high H3K27 binding and lower Prdm3 and Prdm16. CL2 is high Prdm3 and H3K27 with low Prdm16. CL3 is high Prdm16 and H3K27 with low Prdm3. CL4 has high binding of all while CL5 has high Prdm3 only and CL6 is high Prdm3 and Prdm16 only. (E) Heat maps plotting the normalized enrichment of Prdm3, Prdm16, H3K27ac, and IgG relative to +/- 750 bp from the TSS of called peaks. Heatmap is ordered based on identified clusters showing similar enriched peaks as in (D). CL, cluster.

**Figure 2.**
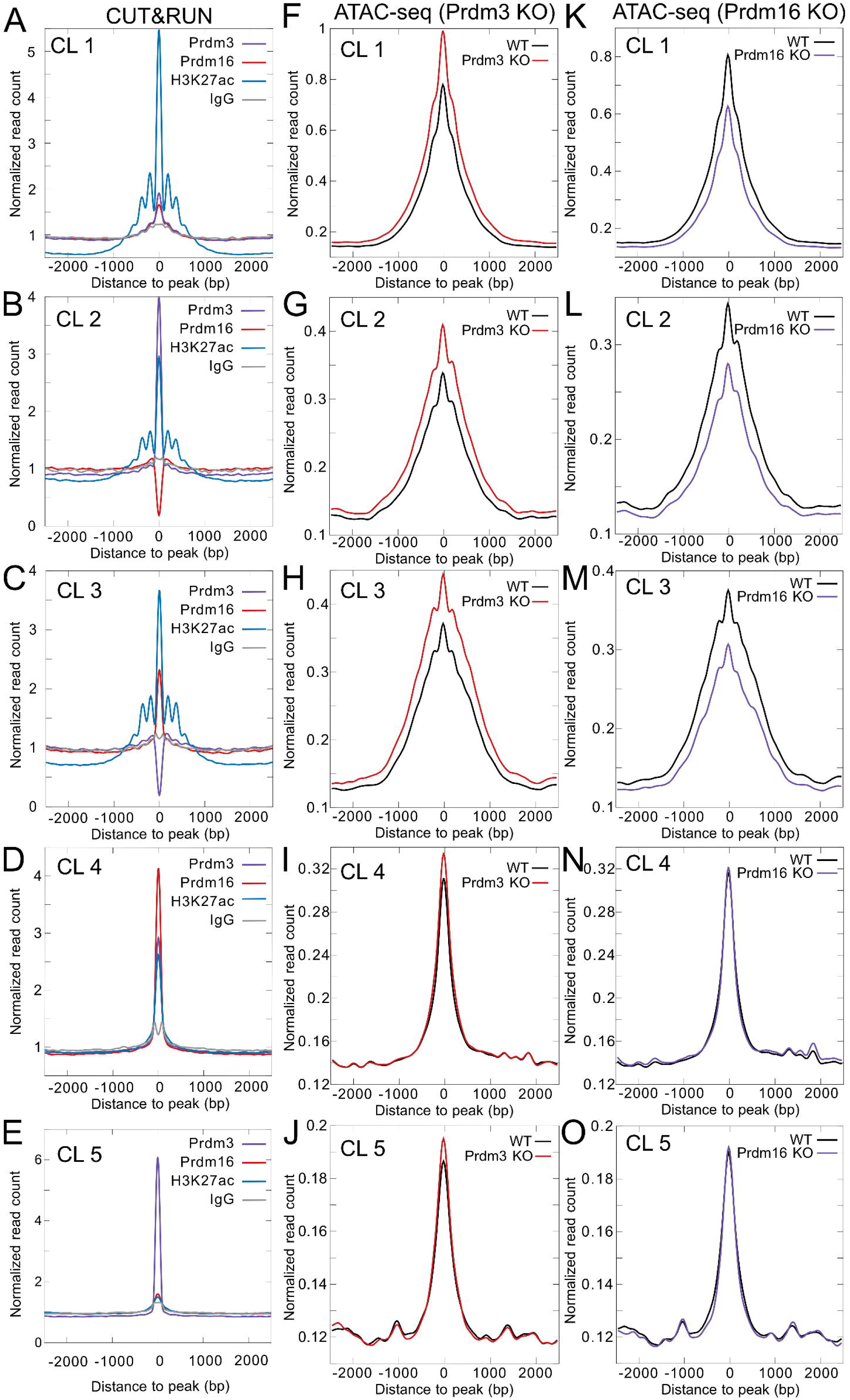
CUT&RUN and ATAC-seq profiles at CUT&RUN peaks. (A-E) Metaplots of normalized read counts +/- 2500 bp relative to peak center for peaks belonging to each of the 5 clusters are shown for CUT&RUN datasets. Each line trace represents the profile of each gene: Purple, Prdm3; Red, Prdm16; Blue, H3K27; and Grey, IgG graphed as normalized read count to distance to peak in base pairs (bp). The H3K27 profiles in CL1-3 (A-C) show a repetitive peaks between 0-1000 that represent the wraps around the nucleosome. Prdm3 and Prdm16 show similar albeit short profile patterns. In CL4-5, there is one strong peak at 0 in all traces. (F-J) ATAC-seq datasets for *prdm3* mutants (*prdm3* KO, red) as compared to wildtype (WT, black) graphed as normalized read count to distance to peak in base pairs (bp). All clusters show traces that are more highly enriched in the *prdm3* mutant then the wildtype. (K-O) ATAC-seq datasets for *prdm16* mutants (*prdm16* KO, blue) as compared to wildtype (WT, black) graphed as normalized read count to distance to peak in base pairs (bp). CL1-3 show traces that have reduced enrichment in the *prdm16* mutant then the wildtype. CL4-5 on the other hand are similar to wildtype.

**Table 1:** Summary of Peak Counts

| Nucleosomal Peaks |  |  |  |
| --- | --- | --- | --- |
| Target | No. of peaks | No. of peaks >2 log2 IgG | % peaks > cutoff |
| IgG | 67,436 |  |  |
| Prdm3 | 119,557 | 36,647 | 31% |
| Prdm16 | 120,162 | 26,863 | 22% |
| H3K27ac | 177,095 | 57,688 | 33% |
| Short Fragment Peaks |  |  |  |
|  | No. of peaks | No. of peaks >2 log2 IgG | % peaks > cutoff |
| IgG | 13,075 |  |  |
| Prdm3 | 24,217 | 606 | 3% |
| Prdm16 | 12,670 | 572 | 5% |
| H3K27ac | 23,956 | 941 | 4% |

Genes associated with each peak were identified using GREAT online server and GO pathway enrichment analysis and WIKI pathways was performed on the nearest gene list associated with each cluster. Putative target genes in Cluster 1(CL1), defined as weaker binding of both Prdm3 and Prdm16, Gene Ontology (GO) and WIKI pathway analysis suggested enrichment and putative target genes in this cluster are associated with stem cell differentiation and cell movement including *itgb5*, *prickle1a*, *cdc42*, and *snail1a* (**Suppl. Figure 1,2**). Cluster 2 (CL2) has only Prdm3 and H3K27ac enrichment at the promoter and is representative of NC and neuronal development as well as canonical Wnt signaling with targets such as *kita*, *sox2*, *foxi1*. Conversely, cluster 3 (CL3) is enriched in Prdm16 and H3K27ac and includes Wnt signaling, pluripotency, NC development/ migration, and axon guidance targets such as *myf5*, *itga1*, *pax3a* and *has1*. Interestingly, both Prdm3 and Prdm16, are highly enriched in Cluster 4 (CL4), with low H3K27ac enrichment, indicating these areas are not actively transcribed but rather in a more poised state. The genes in this cluster are largely associated with neuronal and cranial nerve development, Wnt signaling, iridiphore and include many well-known pigment, neural and NC genes including *dct, dlx3b, foxd3, isl1a, tbx5a, and wnt5a.* Cluster 5 (CL5) contains regions uniquely bound Prdm3 with complete absence of Prdm16 and H3K27ac.

These putative genes include those involved in cell differentiation and canonical Wnt signaling such as *neurog1* and *irx5a*. Cluster 6 (CL6) has minimal enrichment of Prdm3 and Prdm16 without H3K27ac and is represented by nervous system and neurogenesis through GO analysis as well as many aspects of Wnt signaling with genes including *irx5b, mkx1, shh,* and *chd5*.

Together this data indicates that there are both similar and different binding signatures for Prdm3 and Prdm16. These binding signatures suggest both co-regulation of the same genes by both Prdm3 and Prdm16, and also regulation by either Prdm3 or Prdm16 alone.

We next plotted ATAC-seq enrichment at CUT&RUN peaks grouped by the six clusters to compare to the CUT&RUN (**Figure 2**). Cluster 1, which has significant levels of both Prdm3 and Prdm16, but much higher levels of H3K27ac (**Figure 2**), has the highest accessibility across the clusters. Clusters 2 and 3, which have either Prdm3 (Cluster 2) or Prdm16 (Cluster 3) enriched along with high enrichment of H3K27ac show accessibility levels that are half of Cluster 1. Clusters 1-3 show a gain in accessibility upon loss of Prdm3, but a loss of accessibility upon loss of Prdm16 (**Figure 2F-H, K-M**). This suggests that Prdm3 and Prdm16 have opposing functions at these sites: Prdm16 promotes accessibility, whereas Prdm3 promotes compaction. The opposing activities of both these complexes result in the observed steady state accessibility.

Clusters 4 and 5 also have much lower accessibility on average compared to Cluster 1. Clusters 4 and 5, which display either Prdm16 or Prdm3 enrichment predominantly, but are not accompanied by high level of H3K27ac, show minimal changes in accessibility upon loss of either Prdm3 or Prdm16, further suggesting that these sites are at a poised state rather than being actively transcribed.

### *prdm3* and *prdm16* targets have gene expression changes in the mutants

Next, to examine if the targets identified by CUT&RUN were differentially expressed in the mutants, we determined the level of expression in a previous RNA-seq as well as assayed gene expression by HCR in situ hybridization. To do so, we identified genes from each specific cluster that represent a specific GO enrichment pathway and determined if they were differentially expressed in our previously performed RNAseq from Tg*(-4.9sox10:EGFP)* sorted cells at 48 hpf. **Table 2** shows genes from each cluster and their differential gene expression fold change from the RNA-seq between in wildtype and *prdm3* and *prdm16* mutants as well as CUT&RUN score. Consistent with the idea that *prdm3* acts primarily as a repressor with increased expression in the mutants and *prdm16* acts as an activator showing decreased expression in the mutants, we validated these changes in gene expression by HCR analysis for targets from each cluster in wildtype as compared to mutants. Because some of the changes are very low from the RNAseq, we predict we may not observe significant changes by *in situ* hybridization by gene expression analysis. The cluster genes that are examined for expression were chosen based on previous studies suggesting they are expressed in the pharyngeal arches or area of the cranial ganglia at 48 hpf and for the most part, are differentially expressed between *prdm3* and *prdm16* mutants as compared to wildtype controls.

**Table 2.** Selected genes from each cluster and their RNA-seq/CUT&RUN scores

| Gene | RNA-seq |  |  |  | Cluster |
| --- | --- | --- | --- | --- | --- |
|  | PRDM16 log2FC | PRDM16 FDR | PRDM3 log2FC | PRDM3 FDR |  |
| **prickle1a** | -6.45 | 0.113 | 1.19 | 0.70 | CL1 |
| **snai1a** | -1.67 | 0.508 | 1.07 | 0.90 | CL1 |
| **foxi1** | 0 | 99.000 | 1.67 | 0.65 | CL2 |
| **sox2** | -14.64 | 0.063 | 1.8 | 0.32 | CL2 |
| **myf5** | 0 | 0.644 | -5.33 | 99.00 | CL3 |
| **pax3a** | 9.39 | 0.782 | 1.57 | 0.40 | CL3 |
| **foxd3** | -3 | 0.002 | 1.6 | 0.88 | CL4 |
| **isl1** | -2 | 0.011 | 1.56 | 0.37 | CL4 |
| **tbx5a** | -5 | 0.190 | 1.63 | 0.68 | CL4 |
| **wnt5a** | 0 | 0.644 | 3 | 99.00 | CL4 |
| **neurog1** | -1 | 99.000 | 3.36 | 0.57 | CL5 |
| **mkxa** | -4 | 0.023 | -1.35 | 0.80 | CL6 |
| chd5 | 6 | 0.606 | 1.07 | 99.00 | CL6 |

### Gene expression analysis of Cluster 1 targets

For Cluster 1 with the gene ontology around cell movement, we assayed the expression of *prickle1a* and *snail1a* by HCR. *snail1a* is a transcription factor that along with its paralog, *snail2*, is widely known to regulate EMT and migration in many cell types as well as NCCs (LaBonne and Bronner-Fraser, 2000; Sakai et al., 2006; Thisse et al., 1993). *snail1a* is expressed within the sox10+ pharyngeal arches as well as the ventral otic vesicle at 48 hpf in wildtype embryos (**Figure 3**). In *prdm3*-/- embryos, expression is slightly reduced with a modest but insignificant reduction in total corrected total cell fluorescence (CTCF) within the arches and not as well defined as compared to controls. In *prdm16* mutant embryos, expression is observed within the wildtype expression domain and is significantly increased, but not as broadly expressed. *prickle1a* is a member of the PCP signaling pathway and involved in NCC EMT and migration in *Xenopus* and zebrafish (Ahsan et al., 2019; Dingwell and Smith, 2006). Expression of *prickle1a* at later stages (48 hpf) embryos is within the presumed cranial vasculature of the pharyngeal arch region and overlaps with *sox10* expression domain in wildtype (**Suppl. Figure 3**). *prickle1a* expression is spatially comparable between wildtype with reduced CTCF in the pharyngeal arches in *prdm3*-/- while *prdm16* mutants have similar expression in the pharyngeal arch area.

**Figure 3:**
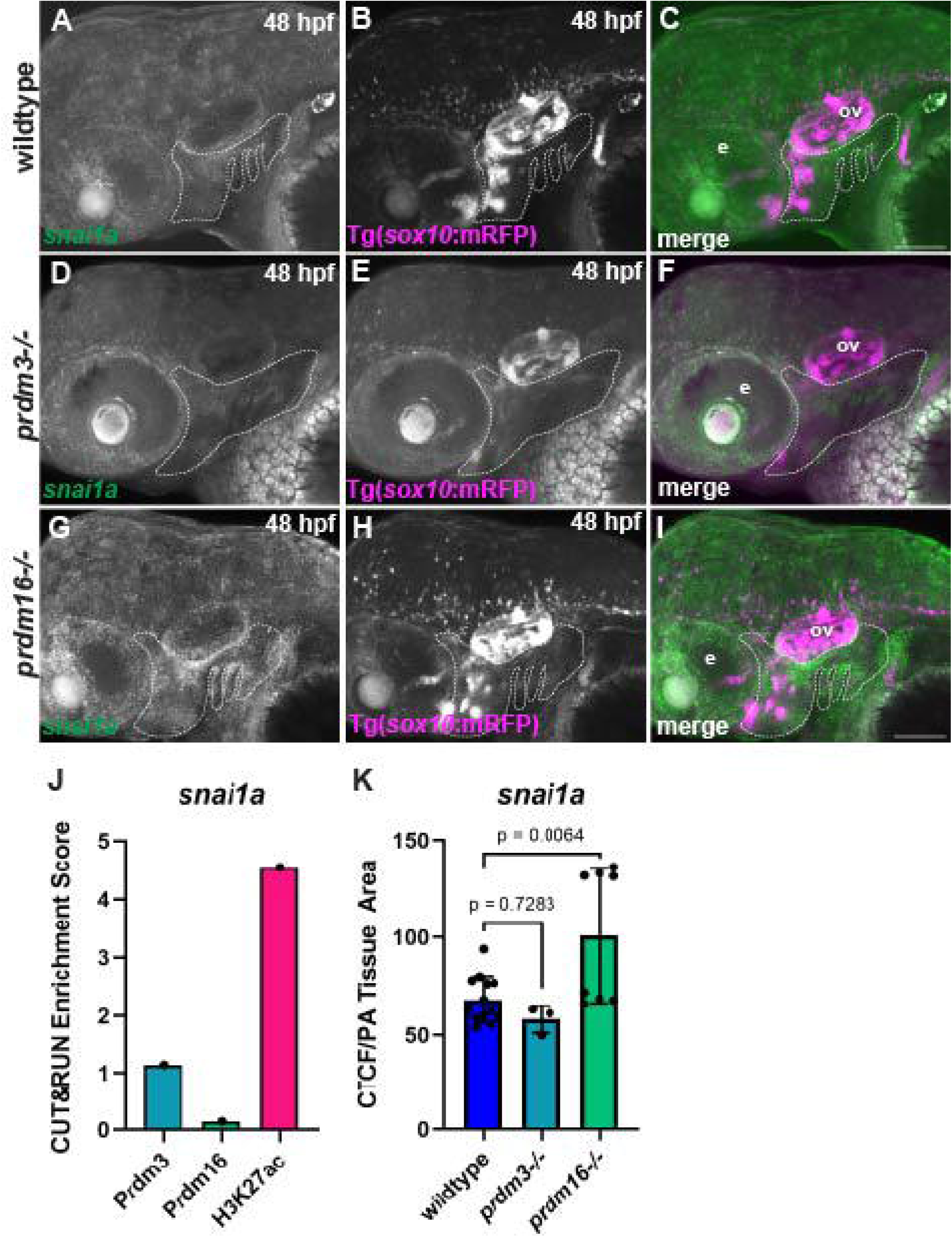
Cluster 1 gene *snail1a* is expressed in the zebrafish pharyngeal arches. Whole-mount HCR in situ hybridization to localize *snail1a* transcripts in zebrafish embryos at 48 hpf in the pharyngeal arches labeled with sox10:mRFP(magenta). Images are lateral views, anterior to the left. (A-C) Representative confocal images of wildtype *Tg(sox10:* mRFP) embryos (magenta) expressing *snail1a* (green) in the pharyngeal arches at 48hpf suggesting weak expression at this time point, n=12. (D-F) Expression of *snail1a* in *prdm3* mutant embryos at 48hpf shows similar expression to wildtype embryos, n=3. (G-I) Expression of *snail1a* in *prdm16*-/- shows a significant increase in fluorescent intensity, n=8. (J) CUT&RUN enrichment score for *snail1a* for Prdm3, Prdm16 and H3K27ac suggesting high H3K27ac binding and low Prdm3 and Prdm16. (K) *snail1a* CTCF controlled for pharyngeal arch (pa) area showing significant increase of expression in *prdm16* mutants. Scale bar is 100um e, eye; ov, otic vesicle. Dash line outlines the pharyngeal arches.

### Gene expression analysis of Cluster 2 targets

Expression of *foxi1* and *sox2* were assessed for Cluster 2 that was enriched in genes important for neural crest, neuron development and Wnt signaling. *foxi1* is an important regulator of preplacodal fate in both zebrafish and mice (Bhat et al., 2012; Edlund et al., 2014; Nechiporuk et al., 2007; Ohyama and Groves, 2004), and functions in the branchial arches (Edlund et al., 2014). By HCR expression analysis we observe similar expression within the arches in *prdm3* mutants with a significantly higher but diffuse area of expression in the *prdm16*-/- embryos (**Figure 4**). *sox2,* a pluripotency transcription factor that is expressed in neural tissue (Kishi et al., 2000; Liu and Labosky, 2008). HCR analysis at 48 hpf shows *sox2* is strongly expressed in the brain, eye, cranial lateral line, the pharyngeal arches and area behind the eye including the otic vesicle. In *prdm3* mutants, *sox2* expression is increased but not significantly by CTCF in the pharyngeal arches but seemingly in the brain. In *prdm16*-/- there is a particularly dramatic decrease of *sox2* expression in the brain but similar expression remains in the pharyngeal arches (**Suppl. Figure 4**).

**Figure 4:**
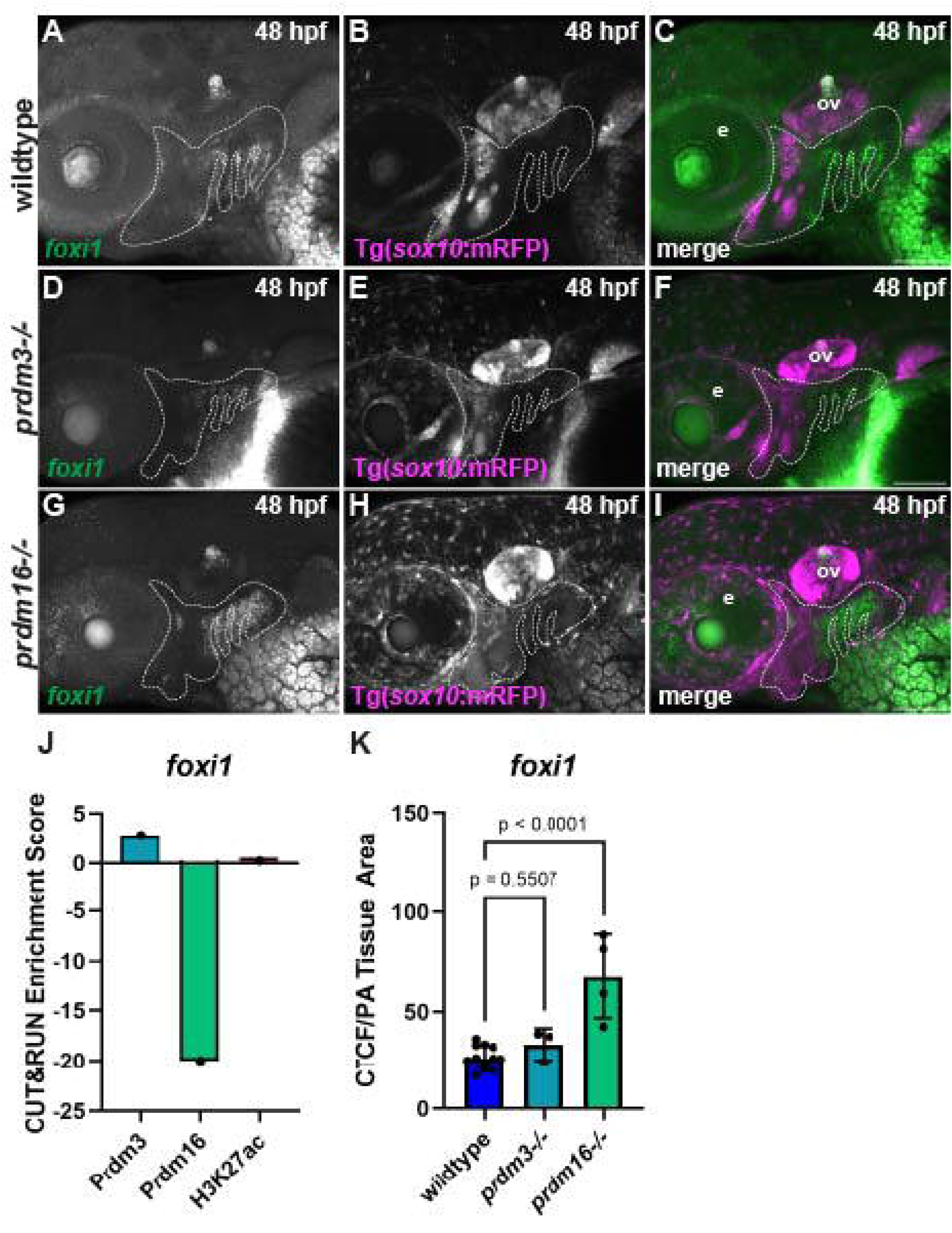
***foxi1* in Cluster 2 is expressed in the zebrafish pharyngeal arches.** Whole-mount HCR in situ hybridization for *foxi1* in zebrafish embryos at 48 hpf in the pharyngeal arches labeled with sox10:mRFP(magenta). Images are lateral views, anterior to the left. (A-C) Wildtype *Tg(sox10:* mRFP) embryo (magenta) expressing *foxi1* (green) in the pharyngeal arches and otic ganglia. In control embryos the expression is in the dorsal pharyngeal arches, n=12. (D-F) Expression of *foxi1* in *prdm3* mutant embryos shows a similar expression as to wildtype controls, n=3. (G-I) Expression of *foxi1* in *prdm16*-/- shows an overall increase and more diffuse area of fluorescent intensity as compared to controls, n=4. (J) CUT&RUN enrichment score for *foxi1* for Prdm3, Prdm16 and H3K27ac showing high Prdm3 enrichment and low Prdm16 and H3K27ac. (K) *foxi1* CTCF controlled for pharyngeal arch (pa) area showing similar expression for *prdm3*-/- and a significant increase of expression in *prdm16* mutants. Scale bar is 100um e, eye; ov, otic vesicle. Dash line outlines the pharyngeal arches.

### Gene expression analysis of Cluster 3 targets

Genes associated with peaks in cluster 3 were enriched in neural crest development and migration, axon guidance, Wnt signaling and pluripotency. *myf5* was primarily expressed in head and trunk muscles as well as the pharyngeal arches. *myf5* expression was reduced in *prdm3* mutants and decreased in the anterior pharyngeal arch domain in *prdm16*-/- (**Figure 5**). *pax3a*, a transcription factor involved in neural plate border, eye and muscle development, is mostly expressed within the brain. By HCR, *pax3a* expression is mostly within the brain at these stages and is slightly but similarly expressed in *prdm3* and *prdm16* mutants (**Suppl. Figure 5**).

**Figure 5:**
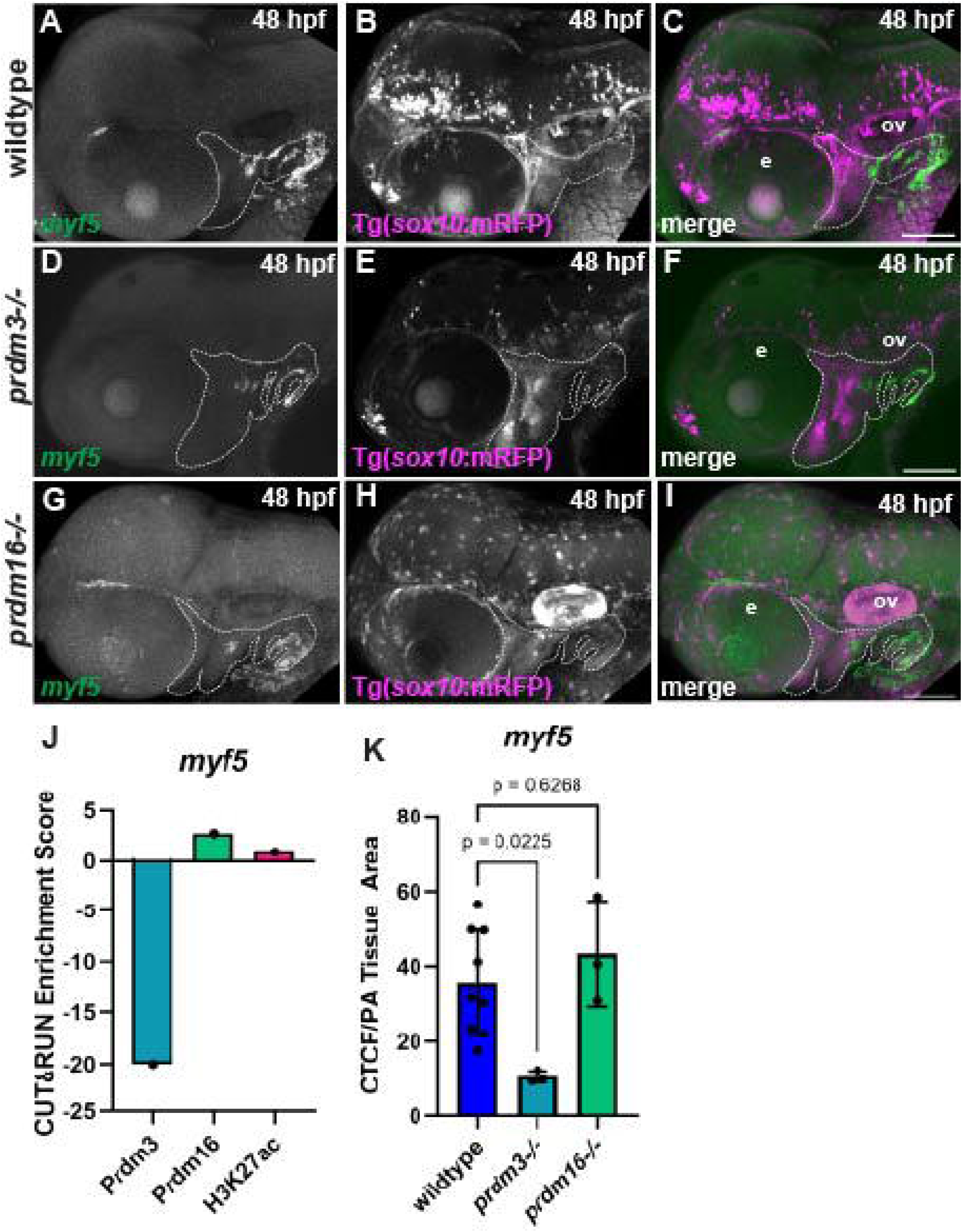
Cluster 3 gene *myf5* is expressed in the zebrafish pharyngeal arches. Whole mount HCR in situ hybridization for *myf5* in zebrafish embryos at 48 hpf in the pharyngeal arch (pa)area. Images are lateral views, anterior to the left. (A-C) Wildtype *Tg(sox10:* mRFP) embryo(magenta) expressing *myf5* (green) in the pharyngeal arches, n=7. (D-F) Expression of *myf5* in *prdm3* mutant embryos shows a trend towards a decrease in overall expression, n=3. (G-I) Expression of *myf5* in *prdm16*-/- shows and overall increase in expression, n=3. (J) CUT&RUN enrichment score for *myf5* for Prdm3, Prdm16 and H3K27ac showing high Prdm16 and low Prdm3 and H3K27ac. (K) *myf5* CTCF controlled for pharyngeal arch (pa) area showing a trending decrease of expression in *prdm3* and similar expression in *prdm16* mutants. Scale bar is 100um e, eye; ov, otic vesicle. Dash line outlines the pharyngeal arches.

### Gene expression analysis of Cluster 4 targets

The genes identified in cluster 4 were enriched in pathways important for neuronal and cranial nerve development, iridophores and Wnt signaling. As such expression of *foxd3* and *isl1a*, were examined for Cluster 4. *foxd3* is a transcription factor that early on is important for NC specification but that later is expressed in pigment and glia fate, and can act as a pioneer factor to both activate and repress transcription (Lukoseviciute et al., 2018; Stewart et al., 2006; Teng et al., 2008). Expression of *foxd3* at this stage 48 hpf is restricted to the CNS and peripheral ganglia glia. In *prdm3* mutants, *foxd3* expression is spatially similar, but reduced to that observed in the wildtype, particularly in the area around the trigeminal ganglia behind the eye and more posterior ganglia in the arch region. In *prdm16*-/- embryos, *foxd3* expression is absent or severely reduced (**Figure 6**).

**Figure 6:**
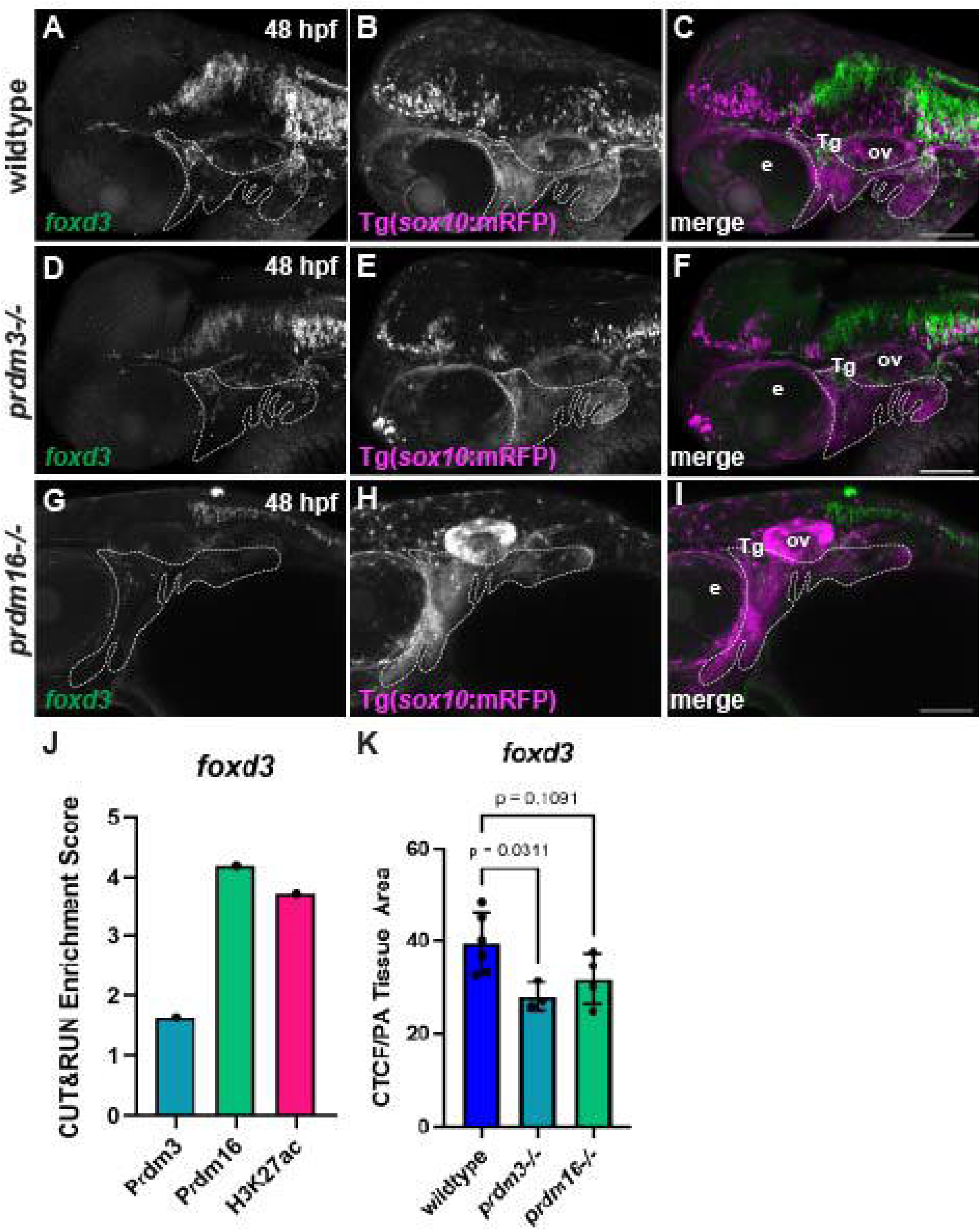
***foxd3* in Cluster 4 is expressed in the area around the trigeminal ganglion in zebrafish.** Whole mount HCR in situ hybridization for *foxd3* in zebrafish embryos at 48 hpf in the pharyngeal arch (pa) area. Images are lateral views, anterior to the left. (A-C) Wildtype *Tg(sox10:* mRFP) embryo (magenta) expressing *foxd3* (green) strongly in the area around the trigeminal ganglia (Tg) and weaker in the pharyngeal arches, n=6. (D-F) Expression of *foxd3* in *prdm3* mutant embryos shows expression levels similar to wildtype controls, n=3. (G-I) Expression of *foxd3* in *prdm16*-/- shows expression levels similar to wildtype controls, n=4. (J) CUT&RUN enrichment score for *foxd3* for Prdm3, Prdm16 and H3K27ac showing high enrichment for all Prdm3, Prdm16 and H3K27ac. (K) *foxd3* CTCF controlled for pharyngeal arch (pa) area showing a similar expression levels in wildtype, *prdm3* and *prdm16* mutants. Scale bar is 100um e, eye; ov, otic vesicle, Tg, trigeminal ganglion. Dash line outlines the pharyngeal arches.

The transcription factor *isl1a* is required for differentiation of motor and sensory neuron populations (Hutchinson and Eisen, 2006; Olesnicky et al., 2010). At 48 hpf *isl1a* expression is localized to the brain and cranial ganglia neurons. In *prdm3* mutants *isl1a* expression by CTCF is similar in the ganglia, with broader and more diffuse domains. Conversely, in *prdm16* mutants, *isl1a* expression is slightly reduced (**Suppl. Figure 6**).

### Gene expression analysis of Cluster 5 targets

Genes identified in cluster 5 were largely associated with cell differentiation and canonical Wnt signaling. Here, we examined *neurog1* expression. *neurog1* is required for specifying cranial ganglia cells (Andermann et al., 2002; Chitnis, 1995; Ma et al., 1998). *neurog1* is a transcription factor that is expressed in the primary sensory neuron populations of zebrafish including cranial ganglia and Rohon-Beard sensory neurons. In later development, it is expressed within the pharyngeal endoderm. In wildtype embryos, HCR shows expression of *neurog1* in the brain, eye and pharyngeal endoderm. In *prdm3*-/- embryos, expression was similarly expressed in the pharyngeal arch endoderm but overall significantly decreased in *prdm16* mutants (**Figure 7**).

**Figure 7:**
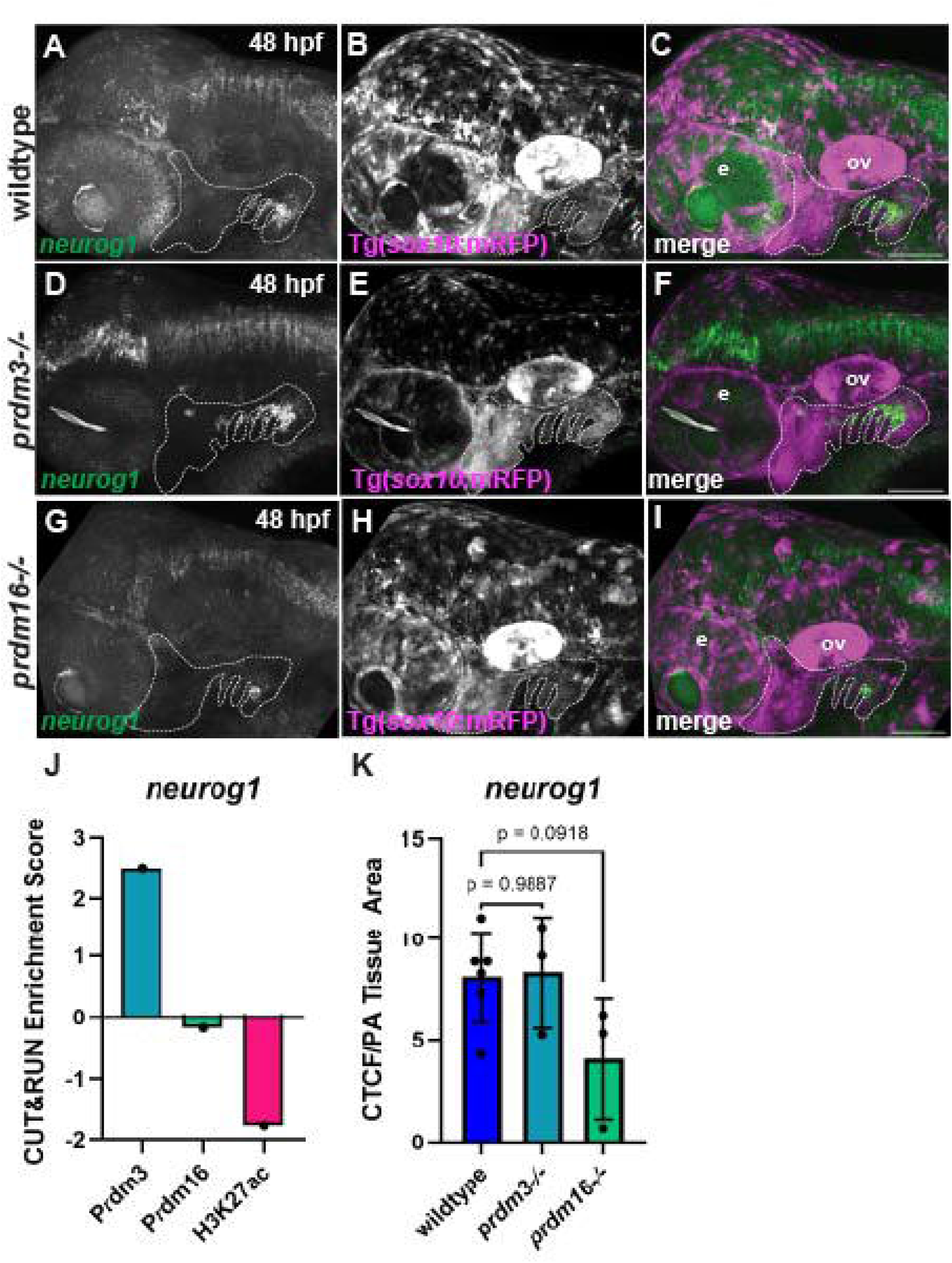
Cluster 5 gene *neurog1* in the zebrafish pharyngeal arches. Whole mount HCR in situ hybridization for *neurog1* in zebrafish embryos at 48 hpf in the pharyngeal arch (pa) area. Images are lateral views, anterior to the left. (A-C) Wildtype *Tg(sox10:* mRFP) embryo (magneta) expressing *neurog1* in the pharyngeal arches, n=6. (D-F) Expression of *neurog1* in *prdm3* mutant embryos showing similar expression to controls, n=3. (G-I) Expression of *neurog1* in *prdm16*-/- show a trending decrease in expression, n=3. (J) CUT&RUN enrichment score for *neurog1* for Prdm3, Prdm16 and H3K27ac showing high Prdm3 and low Prdm16 and H3K27ac. (K) *foxd3* CTCF controlled for pharyngeal arch (pa) area showing a trending decrease of expression in *prdm16* and similar expression in *prdm3* mutants. Scale bar is 100um e, eye; ov, otic vesicle. Dash line outlines the pharyngeal arches.

### Gene expression analysis of Cluster 6 targets

Cluster 6, the smallest cluster, was defined by enrichment of both Prdm3 and Prdm16 without H3K27ac. This cluster is largely characterized by genes associated with neural crest and neurogenesis. Two genes were examined for expression in *prdm3* and *prdm16* mutants, including *mkxa* and *cdh5*. *mkxa* is a homeodomain transcription factor expressed in muscle as well as the pharyngeal arches (Adachi et al., 2022; Chuang et al., 2014). At 48 hpf, *mkxa* is expressed in the first and second arch, the most posterior arch and the otic vesicle. HCR analysis of *mkxa* expression shows a reduction in both mutants, with a significant reduction in the arches of both *prdm3* and *prdm16* mutants (**Figure 8**).

**Figure 8:**
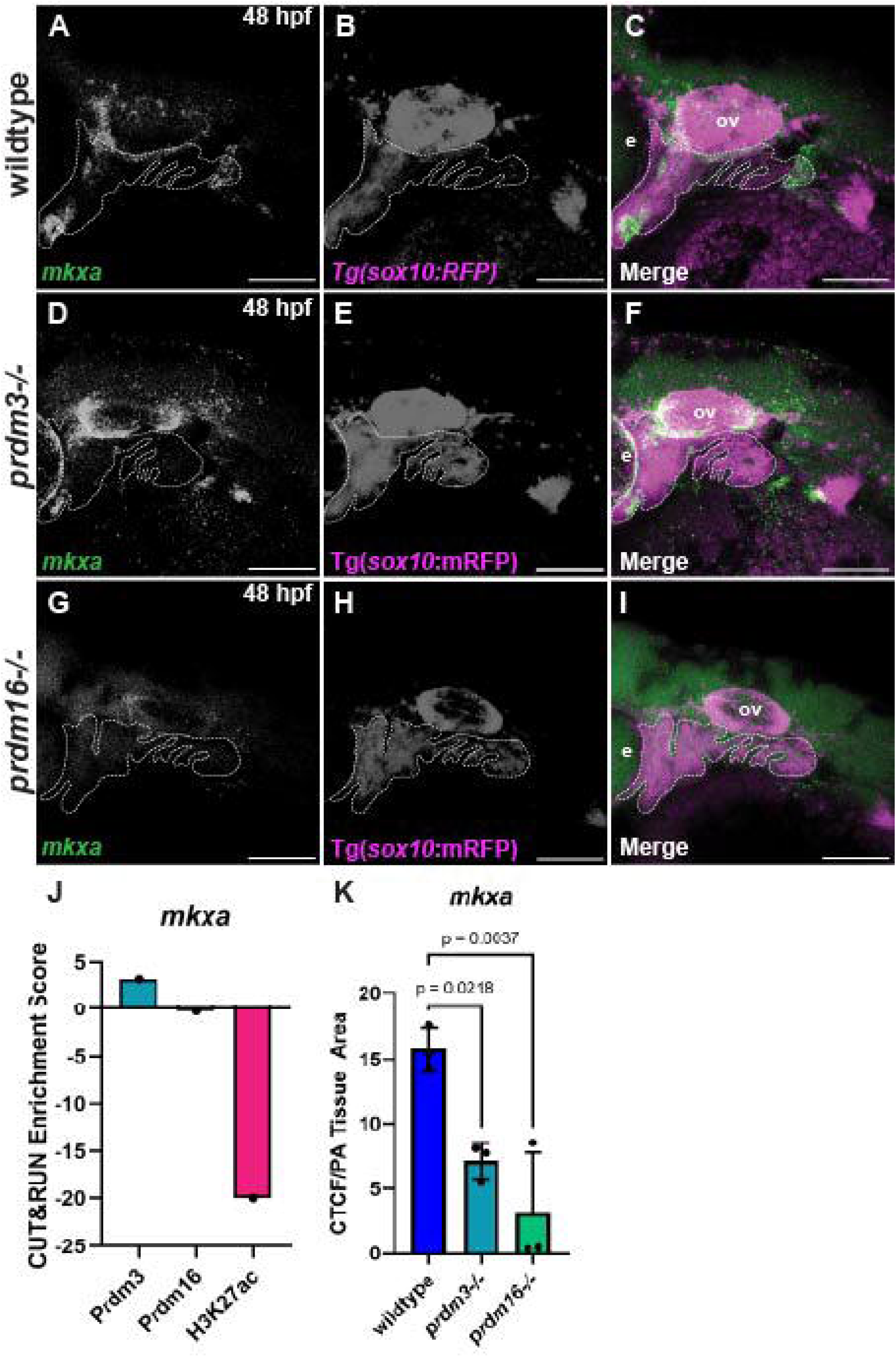
***mkxa* in Cluster 6 is express in the anterior and posterior pharyngeal arches and the area around the otic vesicle.** Whole mount HCR in situ hybridization for *mkxa* in zebrafish embryos at 48 hpf in the area around the pharyngeal arches (pa). Images are lateral views, anterior to the left. (A-C) Wildtype *Tg(sox10:mRFP)* embryo (magenta) expressing *mkxa* (green) in the pharyngeal arches, n=3. (D-F) Expression of *mkxa* in *prdm3* mutant embryos shows a significant reduction in expression, n=3. (G-I) Expression of *mkxa* in *prdm16*-/- shows a significant reduction in expression, n=3. (J) CUT&RUN enrichment score for *mkxa* for Prdm3, Prdm16 and H3K27ac showing high Prdm3 and low Prdm16 and H3K27ac. (K) *mkxa* CTCF controlled for pharyngeal arch (pa) area showing a significant decrease of expression in *prdm3* and *prdm16* as compared to controls. Scale bar is 100um e, eye; ov, otic vesicle. Dash line outlines the pharyngeal arches.

*chd5* is chromatin remodeling protein that is expressed broadly in the head at 48hpf and involved in neuronal differentiation and head development (Bishop et al., 2015; Egan et al., 2013; Hwang et al., 2018; Shrestha et al., 2023). HCR for *chd5* at 48 hpf shows a broad expression pattern in the head and pharyngeal arches. While the overall expression domain is similar, the overall level of expression slightly decreased in both *prdm3* and *prdm16* mutants (**Suppl. Figure 7**).

Overall, the direct targets identified from the Prdm3 and Prdm16 CUT&RUN suggest that these genes identified as differentially expressed are expressed in the correct spatial and temporal pattern similar to *prdm3* and *prdm16* and some altered correspondingly in the respective mutants.

### Prdm3 and Prdm16 enrichment is lost at specific targets in *prdm3* and *prdm16* mutants respectively

Finally, we wanted to determine if PRDM enrichment is lost at putative loci of the downstream target genes. We performed CUT&RUN paired with qPCR with primers flanking predictive binding sites for each putative target gene on whole heads dissected from each respective mutant. Consistent with the binding profiles observed in our CUT&RUN sequencing data, we saw similar enrichment for Prdm3 and Prdm16 patterns for target genes for each cluster. Importantly, we observed decreased enrichment at each predicted target gene with of *prdm3* or *prdm16* (**Figure 9**). For Cluster 1, defined as high H3K27ac and low Prdm3 and Prdm16 enrichment, we designed primers to regions of predicted motifs at *snail1a*. We show high abundance of H3K27ac in wildtype embryos as predicted. Prdm3 binds above background and this binding is reduced in the *prdm3* mutants. For Prdm16, we observe no binding above background. For Cluster 2, which has high Prdm3 and H3K27ac enrichment, we tested binding at *foxi1* and observed elevated levels of Prdm3 that were significantly reduced in *prdm3-/-*. As predicted, there was no significant binding of Prdm16. For Cluster 3, characterized by high Prdm16 and H3K27ac, we assayed *myf5* and as expected we observed that while Prdm3 does not bind, Prdm16 and H3K27ac are strongly enriched and this is significantly decreased in *prdm16* mutants. For Cluster 4 defined by increased abundance of Prdm3, Prdm16 and H3K27ac together, we observed significant enrichment of all three at *foxd3* that is greatly reduced in the respective mutants. Prdm16 overall enrichment is much higher at this locus, which is consistent with the binding profiles for CL4, determined from our CUT&RUN sequencing (**Figure 1**). For Cluster 5, which has binding of Prdm3 without H3K27ac, we examined *neurog1*. Here we observed elevated abundance of Prdm3 in controls that is significantly reduced in *prdm3* mutants. We observed no binding for Prdm16 and H3K27ac. For Cluster 6, we assayed enrichment at *mkxa*. This cluster has binding of Prdm3 and Prdm16 only without H3K27ac and we were unable to see reliable data for this target suggesting an unstable interaction. Overall, these results are consistent with our CUT&RUN sequencing data and show a consistent reduction in the respective mutants suggesting a specific binding interaction at those regions to control the expression of those targets.

**Figure 9:**
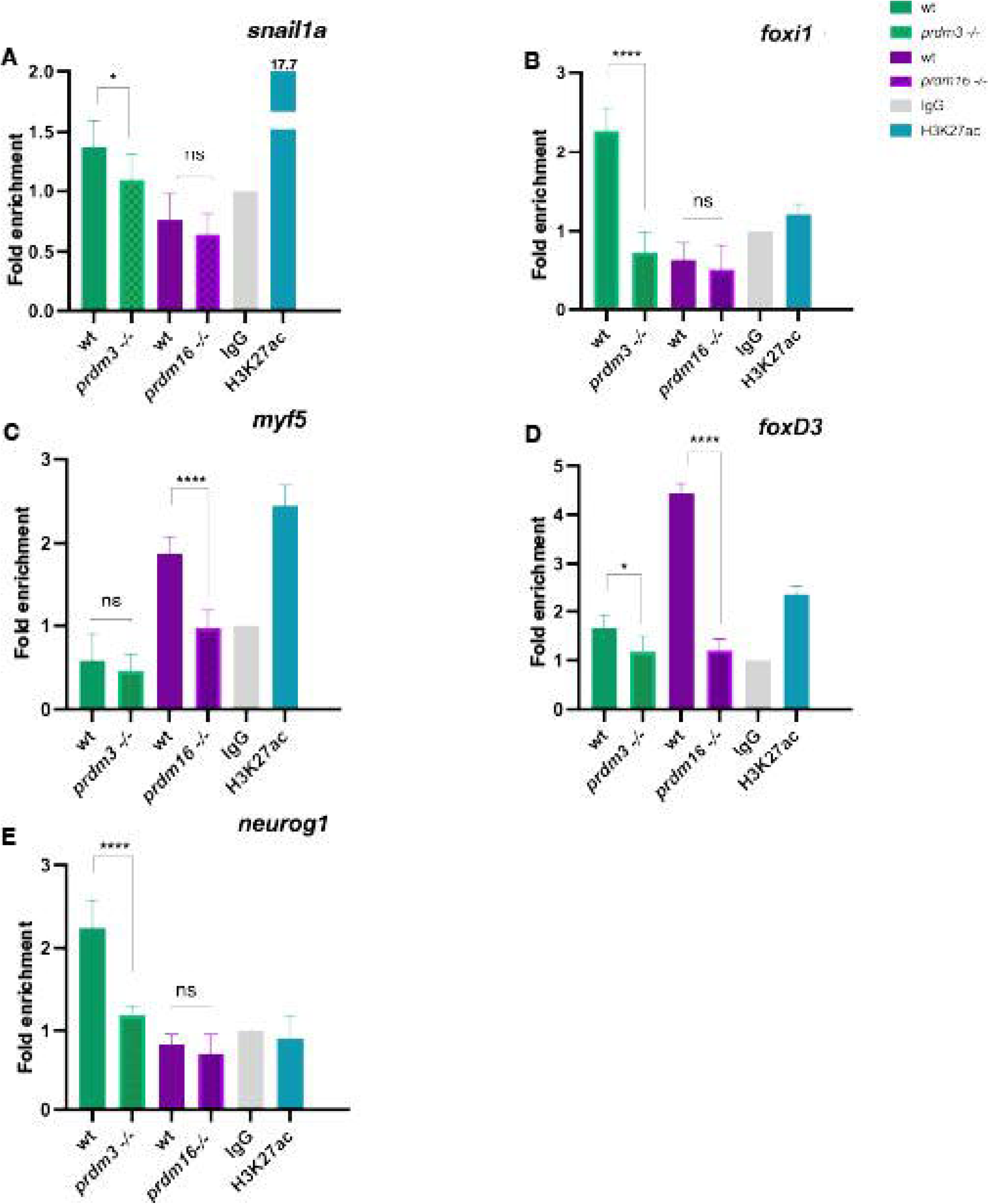
CUT&RUN qPCR identifies direct targets of Prdm3 and Prdm16 that are reduced in *prdm3* and *prdm16* mutants respectively. Graphic illustration comparing wildtype (wt) embryos to *prdm3* and *prdm16* mutants (-/-) showing fold enrichment for each condition. Embryos at 48 hpf were processed for CUT&RUN libraries and qPCR was performed for each target with Prdm3 and Prdm16 in wildtype and mutant embryos and IgG and H3K27ac in wildtype embryos. Three replicates for each genomic loci were performed. (A) *snail1a* enrichment is slightly significantly lower in *prdm3-/-* embryos and similar in *prdm16* mutant embryos with high enrichment of H3K27ac at this locus as compared to controls (B) *foxi1* enrichment is significantly lower in *prdm3-/-* embryos and similar in *prdm16* mutant embryos with no enrichment of H3K27ac at this locus as compared to controls (C) *myf5* enrichment is similar in *prdm3-/-* embryos and significantly lower *prdm16* mutant embryos with high enrichment of H3K27ac at this locus as compared to controls (D) *foxd3* enrichment is significantly lower in both *prdm3-/-* and *prdm16* mutant embryos with high enrichment of H3K27ac at this locus as compared to controls (E) *neurog1* enrichment is significantly lower in in *prdm3-/-* embryos and similar in *prdm16* mutant embryos with no enrichment of H3K27ac at this locus as compared to controls.

## Discussion

In this study, we investigated transcription factor modality of two important regulators of craniofacial development, Prdm3 and Prdm16 by CUT&RUN analysis. We show that there is differential binding activity of the two factors, in relation to H3K27ac, into discrete clusters, that changes in gene expression of Prdm3 and Prdm16 putative target genes correlates with their binding to regions of these genes, and that these interactions are reduced in the mutants. These data support the model that Prdm3 and Prdm16 have dynamic binding ability in which they can bind together at specific targets as well as separately, and either with or without H3K27ac in order to regulate various aspects of cNCC cell fate. These include cell movement during migration, Wnt signaling pathways, as well as different levels of NCC differentiation into neurons, pigment cells and other cellular processes involved in NCC biology.

CUT&RUN sequencing is a relatively new technique to determine transcription factor interactions with DNA, similar to chromatin immunoprecipitation, that instead utilizes enzymatic digestion around DNA bound fragments (Skene and Henikoff, 2017). Unlike chromatin immunoprecipitation, CUT&RUN preserves the fragment size distribution generated during cleavage allowing us to quantify the differences in large fragments ∼100 bases vs short fragments that represent binding associated with nucleosomes and direct transcription factor binding respectively (Tonsager et al., 2025; Trouth et al., 2024). From our analysis, we found Prdm3 and Prdm16 have a preferred nucleosome enrichment of larger fragments from their CUT&RUN datasets compared to the IgG control dataset. Given the fact that both these factors have DNA binding domains as well as SET domains with methyltransferase activity, it is interesting that they predominantly prefer to associate with nucleosomes. Future experiments will determine the protein binding partners required, including both activator and repressor complexes.

There has been an increased interest in the role of PRDM proteins in cell fate determination. Many studies have defined the role for PRDM’s in regulating important cell fate decisions at critical cell lineage choice points. Recent work in the mouse lung suggests that in PRDM3/16 knockout mice, alveolar type 2 (AT2) cells are lost, which are required for surfactant production while gaining alveolar type 1 cells (He et al., 2024). The authors further show that together with NKX2.1, PRDM3/16 regulate the specific differentiation of AT2 cell fate by regulating chromatin accessibility at NKX2.1 targets. Similarly, Prdm1, another family member has been shown to interact with FoxA transcription factors and the nucleosome remodeling complex NuRD to regulate endoderm differentiation (Matsui et al., 2025). Our previous work has shown that Prdm3 and Prdm16 regulates cranial NCC differentiation into chondrocytes, by promoting the differentiation of these cells while simultaneously repressing other cranial NCC cell fates (Shull et al., 2022; Shull et al., 2020). We hypothesize that cartilage/neuron/glia progenitor cell is the likely target of Prdm3 and Prdm16 activity, acting primarily as repressors of transcription of the alternative cell fate. This is consistent with the observation that many of the targets we have identified here by CUT&RUN are neuronal, and that this fate is repressed by Prdm3 and Prdm16 as they activate cartilage and bone. However, given that there are a substantial number of downregulated genes in *prdm3* and *prdm16* mutants at 48hpf, it is possible that Prdm proteins may act as activators at some targets as well. Examination of putative target gene expression in the craniofacial region confirms that many of the *prdm3* target genes generally have an increased expression in the mutants, suggesting Prdm3 normally functions as a repressor. On the other hand, loss of *prdm16* results in mostly downregulated gene expression. The potential activator activity of Prdm16 may not be direct as it could be that the role of *prdm16* is to repress repressors and thereby activate gene expression. Further experiments are required to test this idea.

## Conclusion

In summary, we present here data that supports the role of Prdm3 and Prdm16 associating with nucleosomes for DNA binding. The direct targets identified are generally expressed in the pharyngeal arches and are altered in the respective mutants when Prdm3 and Prdm16 binding is reduced. Overall, these results support a role for Prdm3 and Prdm16 in regulating cell fate determination in cNCCs consistent with their role in other systems and the broader landscape of defining progenitor programs.

## Acknowledgements

We thank the past and present members of the Artinger lab for project feedback, Christine Archer at the University of Colorado Anschutz, Jennifer Sharp at the University of New Mexico, and Marc Tye at the University of Minnesota and the zebrafish fish facility teams for excellent animal care. This work is supported by K99/R00 DE0224034 to L.C.S. and NIDCR R01DE030377 to K.B.A.

---

**Supplemental Figure 1.** : GO analysis of each cluster. Gene Ontology (GO) pathway analysis suggested enrichment and putative target genes in each cluster. (A) Cluster 1 are associated with stem cell differentiation and cell movement. (B) Cluster 2 is representative of neural crest and neuronal development as well as canonical Wnt signaling. (C) Cluster 3 (CL3) is includes Wnt signaling, pluripotency, neural crest development/ migration. (D) Cluster 4 are largely associated with neuronal and cranial nerve development, Wnt signaling, iridiphore and include many well-known pigment, neural and neural crest genes. (E) Cluster 5 include targets involved in cell differentiation and canonical Wnt signaling. (F) Cluster 6 is represented by nervous system and neurogenesis through GO analysis as well as many aspects of Wnt signaling.

**Supplemental Figure 2:** WIKI pathway analysis. WIKI pathway analysis suggested enrichment and putative target genes in each cluster. (A) Cluster 1 are associated mRNA processing, cell adhesion and metabolic signaling. (B) Cluster 2 is representative of neural crest and as well as canonical Wnt signaling. (C) Cluster 3 (CL3) includes Wnt signaling, pluripotency, neural crest development/ migration. (D) Cluster 4 are largely associated with neural crest development, Wnt signaling and pluripotency. (E) Cluster 5 include targets involved in various signaling pathways as well as canonical Wnt signaling. (F) Cluster 6 is represented by many aspect of the Wnt signaling pathway as well as others.

**Supplemental Figure 3.** : Cluster 1 gene *prickle1a* is expressed in zebrafish vasculature. Whole mount HCR in situ hybridization for *prickle1a* in zebrafish embryos at 48 hpf in the pharyngeal arch area. Images are lateral views, anterior to the left. (A-C) Wildtype *Tg(sox10:* mRFP) embryo (magenta) expressing *prickle1a* (green) in the pharyngeal arches, n=8. (D-F) Expression of *prickle1a* in *prdm3* mutant embryos shows a trending reduction in expression, n=3. (G-I) Expression of *prickle1a* in *prdm16*-/- showing similar expression to controls, n=3. (J) CUT&RUN enrichment score for *prickle1a* for Prdm3, Prdm16 and H3K27ac showing high H3K27ac low Prdm3 and low Prdm16. (K) *prickle1a* CTCF controlled for pharyngeal arch (pa) area showing a treading decrease of expression in *prdm3-/-* and a similar level in *prdm16* mutants as compared to controls. Scale bar is 100um e, eye; ov, otic vesicle. Dashed line indicates the pharyngeal arches.

**Supplemental Figure 4.** : *sox2* in cluster 2 is expressed in the zebrafish pharyngeal arches. Whole mount HCR in situ hybridization for *sox2* in zebrafish embryos at 48 hpf in the pharyngeal arch area. Images are lateral views, anterior to the left. (A-C) Wildtype *Tg(-4.9sox10:* mRFP) embryo (magenta) expressing *sox2* (green) in the pharyngeal arches, n=4. (D-F) Expression of *sox2* in *prdm3* mutant embryos shows domains of increased expression, n=4. (G-I) Expression of *sox2* in *prdm16*-/- shows domains of increased expression, n=3. (J) CUT&RUN enrichment score for *sox2* for Prdm3, Prdm16 and H3K27ac showing high Prdm3 and H3K27ac and low Prdm16. (K) *sox2* CTCF controlled for pharyngeal arch (pa) area showing a treading increase of expression in both *prdm3-/-* and in *prdm16* mutants as compared to controls. Scale bar is 100um e, eye; ov, otic vesicle. Dashed line outlines the pharyngeal arches

**Supplemental Figure 5.** : *pax3a* from cluster 3 is expressed in the zebrafish brain. Whole mount HCR in situ hybridization for *pax3a* in zebrafish embryos at 48 hpf in the pharyngeal arch area. Images are lateral views, anterior to the left. (A-C) Wildtype *Tg(sox10:* mRFP) embryo (magenta) expressing *pax3a* (green) primarily in the brain and weakly in the pharyngeal arches, n=9. (D-F) Expression of *pax3a* in *prdm3* mutant embryos show a trending decrease in expression, n=3. (G-I) Expression of *pax3a* in *prdm16*-/- is similar as compared to controls, n=3. (J) CUT&RUN enrichment score for *pax3* for Prdm3, Prdm16 and H3K27ac showing enrichment of Prdm 16 and H3K27ac and low Prdm3. (K) *pax3a* CTCF controlled for pharyngeal arch (pa) area showing a treading decrease of expression in *prdm3-/-* and a similar level in *prdm16* mutants as compared to controls. Scale bar is 100um e, eye; ov, otic vesicle. Dashed line outlines the pharyngeal arches.

**Supplemental Figure 6.** : *isl1*a in cluster 4 is expressed in the zebrafish brain, pharyngeal arches and trigeminal ganglion. Whole mount HCR in situ hybridization for *isl1a* in zebrafish embryos at 48 hpf in the area of the pharyngeal arches. Images are lateral views, anterior to the left. (A-C) Wildtype *Tg(sox10:* mRFP) embryo (magenta) expressing *isl1a* (green) in the pharyngeal arches and trigeminal ganglion, n=7. (D-F) Expression of *isl1a* in *prdm3* mutant embryos is similar to that of controls, n=5. (G-I) Expression of *isl1a* in *prdm16*-/- is trending lower as compared to controls, n=4. (J) CUT&RUN enrichment score for *isl1a* for Prdm3, Prdm16 and H3K27ac showing enrichment for all, Prdm3, Prdm16 and H3K27ac. (K) *isl1a* CTCF controlled for pharyngeal arch (pa) area showing a treading decrease of expression in *prdm16-/-* and a similar level in *prdm3* mutants as compared to controls. Scale bar is 100um e, eye; ov, otic vesicle. Dashed line outlines the pharyngeal arches.

**Supplemental Figure 7.** : *chd5* from cluster 6 is expressed in the zebrafish pharyngeal arches. Whole mount HCR in situ hybridization for *chd5* in zebrafish embryos at 48 hpf in the area of the pharyngeal arches. Images are lateral views, anterior to the left. (A-C) Wildtype *Tg(-4.9sox10:EGFP)* embryo (magenta) expressing *chd5* (green) diffusely in the pharyngeal arches, n=5. (D-F) Expression of *chd5* in *prdm3* mutant embryos shows a slight reduction in expression, n=3. (G-I) Expression of *chd5* in *prdm16*-/- shows a slight reduction in expression, n=4. (J) CUT&RUN enrichment score for *chd5* for Prdm3, Prdm16 and H3K27ac showing enrichment for Prdm3 only and now Prdm16 and H3K27ac. (K) *chd5* CTCF controlled for pharyngeal arch (pa) area showing a treading decrease of expression in both *prdm3* and *prdm16* mutants. Scale bar is 100um e, eye; ov, otic vesicle. Dashed line outlines the pharyngeal arches.

**Supplemental Table 1.**
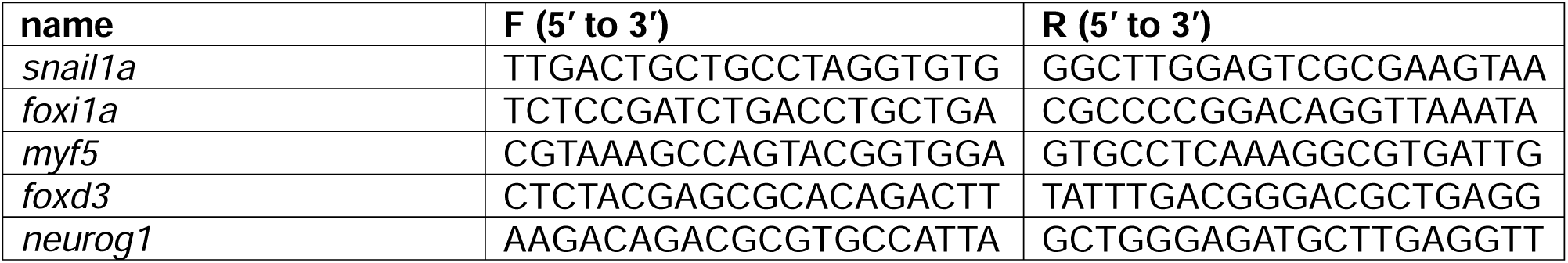
CUT&RUN Primer sequences

