## Supplementary figures and images for "PRDM3 and PRDM16 define cranial neural crest cell states in zebrafish development"

### Supplemental figure1

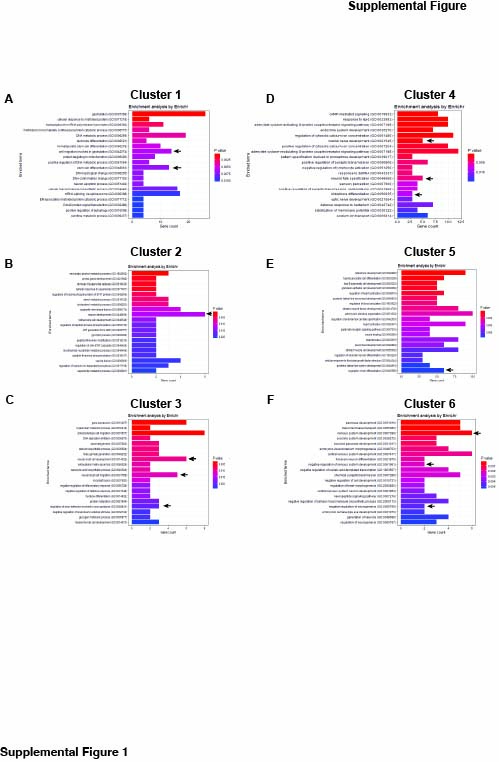

### Supplemental figure2

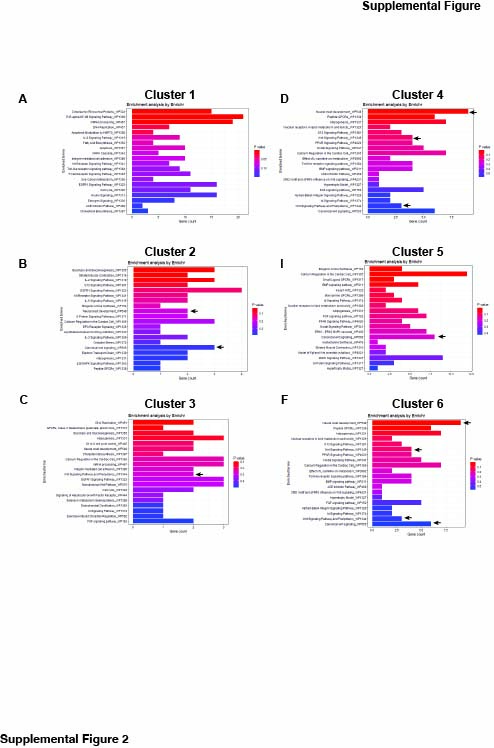

### Supplemental figure3

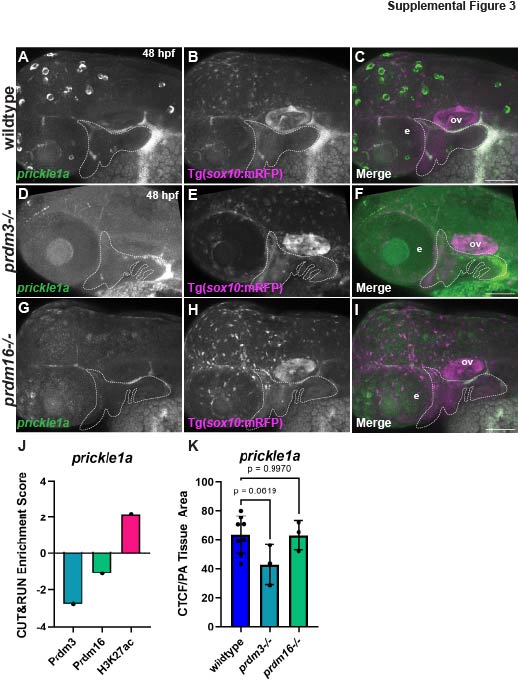

### Supplemental figure4

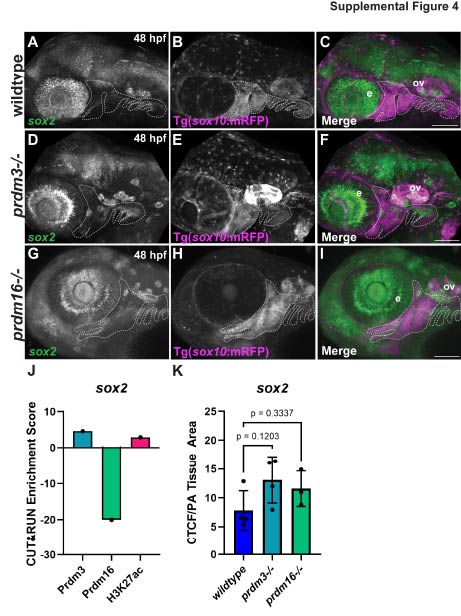

### Supplemental figure5

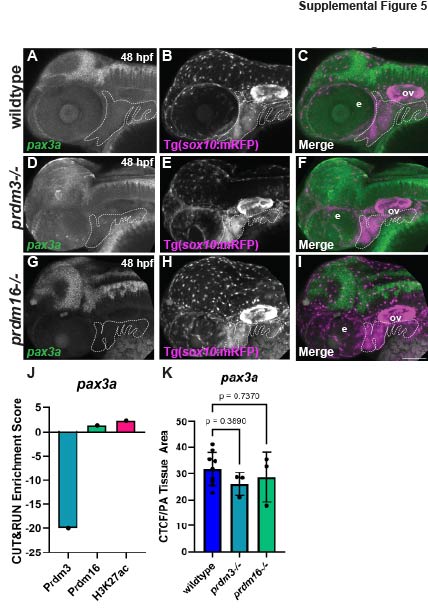

### Supplemental figure6

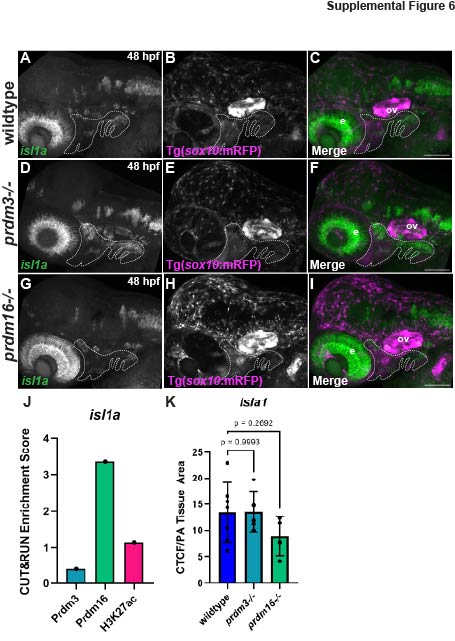

### Supplemental figure7

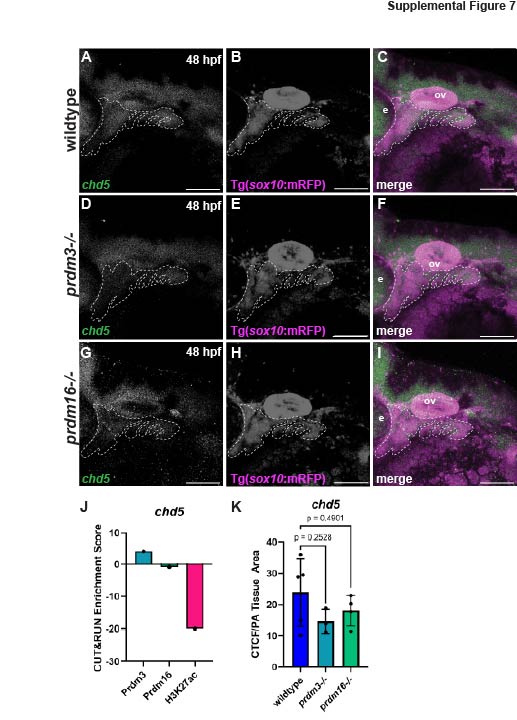
